# Weighted sliced inverse regression for scalable supervised dimensionality reduction of spatial transcriptomics data

**DOI:** 10.1101/2025.07.08.663651

**Authors:** Maximilian Woollard, Pratibha Panwar, Luke T.G. Harland, Jeremy Mo, Hanyun Zhang, Alexander Swarbrick, Berthold Gottgens, Shila Ghazanfar, Linh H. Nghiem

## Abstract

With the dramatic take-up of spatially resolved transcriptomics biotechnologies, performing spatially-aware analysis of the resulting data is crucial to maximise advances in biological understanding. Dimensionality reduction is a first step in almost any analysis of spatial transcriptomics data, regardless of whether the data is collected at the single-cell or spot level. While common approaches, such as principal component analysis, aim to identify low-dimensional scores that preserve total variances of gene expression features, such variances do not usually correspond to biologically relevant spatial variation. To this end, we have developed weighted sliced inverse regression (wSIR), a sufficient dimension reduction technique that performs dimensionality reduction and retains as much predictive power of the spatial coordinates as possible. As a linear dimensionality reduction approach, wSIR is applicable to multiple distinct spatial transcriptomic datasets, and is extremely scalable due to the algorithm and our Rcpp implementation, with over 100, 000 cells processed in under 2 minutes on a standard laptop. The feature loadings are interpretable, and new non-spatial data can be projected into the wSIR low-dimensional space for further downstream analyses. We examine wSIR’s performance through benchmarking and demonstrate its capability of biological discovery through two case studies in breast cancer and early embryonic development.

## 1 Introduction

Spatial transcriptomics is becoming a widespread method for profiling hundreds to thousands of genes at single cell resolution in their spatial context for a variety of tissues (You et al., 2024). There is great potential to uncover the complexity of tissue organisation and discover new biological aspects in normal processes and the emergence of disease. Current analysis of spatial transcriptomics data has emerged primarily as an extension of the last 5-8 years of single-cell transcriptomics research (https://www.scrna-tools.org/), with several key analytical steps often borrowed from single-cell analysis, including normalisation, filtering and dimensionality reduction techniques. Dimensionality reduction, in particular, is a crucial step in analysing complex high-dimensional data, including single-cell and spatial transcriptomics, as several other downstream analysis steps rely on this (Huang et al., 2022; Wang et al., 2023; Cantini et al., 2021). For example, clustering of cells is typically performed using graph-based approaches on low-dimensional embeddings rather than the original gene expression matrix to avoid the curse of dimensionality. As such, effective and appropriate dimensionality reduction of spatial transcriptomics data is an important challenge in the analysis of these data.

Despite the rapidly accelerating generation of spatial transcriptomics data, it is not straightforward how to incorporate spatial coordinates and gene expression measurements to form a low-dimensional representation of the data. Popular dimensionality reduction approaches such as principal component analysis (PCA) do not utilise spatial coordinates; it focuses on finding orthogonal linear combinations of the gene expression data to preserve the largest amounts of their total variation. Other classical statistical approaches can be adapted to perform a “spatially-aware” dimensionality reduction, such as partial least squares (PLS) and linear discriminant analysis (LDA). PLS treats spatial coordinates in each dimension as separate responses, and may not be well suited to account for the complex spatial associations between the spatial coordinates and the gene expressions. LDA is sensitive to the arbitrary choice of allocation of the spatial coordinates into distinct classes. Given these challenges, new approaches have emerged that aim to integrate gene expression and spatial coordinates to arrive at meaningful dimensionality reduction. Some innovative approaches, such as GraphPCA (Yang et al., 2024), SpatialPCA (Shang and Zhou, 2022), MUSTARD (Zhuang et al., 2024), and DR-SC (Liu et al., 2022), improve PCA by incorporating an additional step, such as augmenting the PCA objective function with a spatial penalty term, including a cell-cell kernel function, utilising experimental trajectory information, or incorporating PCA into a hierarchical model, respectively. Other approaches have adapted different modelling strategies, such as nonnegative matrix factorisation, to utilise spatial information (Wang et al., 2024; Georgaka et al., 2023; Townes and Engelhardt, 2022). In another line of research, several methods have utilised a dimensionality reduction step to cluster cells or spots in a “spatially-aware” manner, including STAMP (Zhong et al., 2024), stLearn (Pham et al., 2020), and FAST (Liu et al., 2023). Other approaches used bespoke techniques utilising deep learning to learn from spatial gene expression and images jointly (Tang et al., 2024), or exploited higher order interactions to identify clusters of interest (Haviv et al., 2024). Additionally, some methods are developed to utilise the spatial transcriptomic data as a reference to predict the latent spatial coordinates of scRNA-seq derived data, such as CellContrast (Li et al., 2023) and Tangram (Biancalani et al., 2021).

In statistics research, advances in sufficient dimension reduction (SDR) have emerged in the last few decades. SDR is a technique that enables dimensionality reduction of features so that the ability to predict the response variables is best preserved. Among SDR methods, sliced inverse regression (SIR), which was originally proposed in the seminal paper of Li (1991) is still the most popular due to its simplicity, flexibility, and computational efficiency (Li, 2018). Despite many developments of SIR to handle complex settings, to the best of our knowledge, SIR has not been adapted to be suitable for single-cell or spatial transcriptomics data. While several approaches have been developed for spatial dimensionality reduction, many are prohibitive in terms of computational scalability and runtime, and are limited in their ability to project new gene expression data into a low-dimensional space.

Given these challenges, we have developed <u>w</u>eighted <u>S</u>liced <u>I</u>nverse <u>R</u>egression (wSIR), a new statistical approach that performs SDR to preserve the association between the gene expressions and the spatial information. wSIR takes into account the slice-slice correlation induced from spatial transcriptomics data, and enables projection of non-spatial gene expression data into the low-dimensional spaces. Due to its linearity, wSIR is highly scalable and can be applied to massive single-cell spatial transcriptomics datasets across multiple distinct samples. In this paper, we describe the wSIR approach, explore its properties, and demonstrate its superior performance compared to other statistical methods. Furthermore, we will also employ wSIR to uncover additional insights from spatial transcriptomics data in two real biological contexts in human breast cancer and mouse development.

## 2 Results

### 2.1 Spatially-aware dimensionality reduction with weighted sliced inverse regression (wSIR)

Given the challenges in effectively describing spatial transcriptomics data, we developed a new linear sufficient dimension reduction approach named weighted sliced inverse regression (wSIR). Assuming a *n × p* gene expression matrix **X** with *n* samples and *p* features, and a corresponding *n × q* responses **Y**, sufficient dimension reduction is a class of statistical techniques that aims to estimate a *d* ≪ *p* vector-valued function of the data *f* (**X**) so that **Y** ⊥ **X** | **X** *f* (**X**), that is, that **Y** and **X** are conditionally independent given the reduced data *f* (**X**). Sliced inverse regression (SIR), which was invented in Li (1991) is an example of linear sufficient dimension reduction, where *f* (**X**) is assumed to be a linear combination of the **X**, i.e *f* (**X**) = **XB** with **B** a *p × d* matrix. Under mild technical conditions, SIR has been shown to be a powerful technique in the area of linear SDR due to its ability to preserve complex and nonlinear patterns within the data (Li, 2018; Lin et al., 2019; Girard et al., 2022, among many others). One important challenge in implementing such an approach to spatial transcriptomics, however, is that SIR does not account for possible spatial relationships among the slices, prompting our development of a novel weight component to SIR to create wSIR (Methods).

Briefly, wSIR takes in spatial transcriptomics data with appropriately normalised, centred and scaled gene expression values, alongside spatial coordinates that can belong to one or more distinct tissues or samples (Figure 1a). The observations within each sample are then grouped according to their proximity in physical space into distinct “slices”, alongside the corresponding gene expression data (Figure 1b). Then, a slice-slice correlation matrix is estimated by taking into account the physical proximity of the centroids of each slice per sample (Figure 1c, Methods), where the closer the two slices are, the higher the corresponding value in the correlation matrix. On each slice, gene expression values are averaged over each gene. Following this step, the slice-mean gene expression and slice-slice correlation matrix are subjected to an eigendecomposition to obtain the eigenvectors, which are finally multiplied by the inverse of the sample covariance matrix of the gene expression to the wSIR direction (loadings). A sample-level scores (embeddings) matrix is then formed by multiplying the original data by the loadings matrix, representing the low-dimensional embedding of the data (Figure 1d). This procedure enables the projection of new, unseen, potentially non-spatial data to the same low-dimensional embedding via matrix multiplication. Overall, wSIR enables interpretation of important genes via examination of the loadings matrix, as well as further downstream analysis (such as clustering) on the spatial transcriptomic data, alongside mapping of non-spatial (e.g. scRNA-seq resolved) transcriptomic data to a spatially-informed low dimensional space.

**Figure 1:**
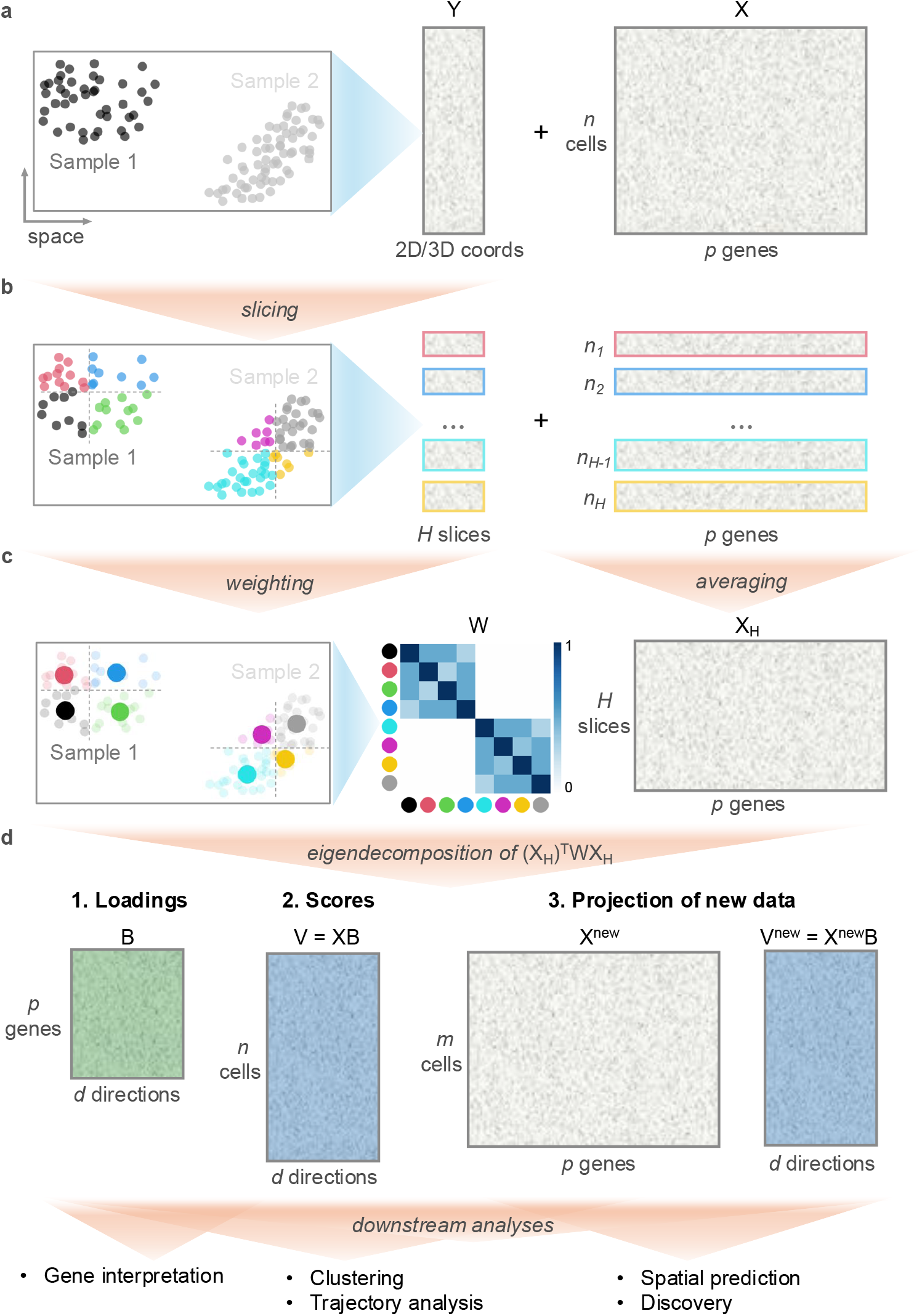
Schematic of wSIR algorithm. **a**, wSIR requires an input of a set of 2D or 3D spatial coordinates for cell-resolved for spot-resolved spatial omics, alongside a normalised and centred gene expression matrix, potentially coming from multiple distinct samples. **b**, wSIR performs slicing of the spatial coordinates per sample, and groups cells/spots into slices according to their proximity in space. **c**, A slice-slice similarity matrix is created where entries correspond to the proximity of slice centroids in space. Slices belonging to different samples are assigned a value of zero, indicating no proximity. The corresponding sets of cells/spots are summarised by taking averages across all of the genes. **d**, wSIR then performs eigendecomposition of the slice-summarised gene expression data and slice-slice similarity. This provides three main outputs, 1. loadings, 2. scores for the input data, and 3. an ability to project new gene expression data into the same low-dimensional space. Overall, wSIR enables downstream analyses for biological discovery.

### 2.2 wSIR prioritises spatially relevant variation

To examine the properties of wSIR, we created a simple simulation containing features that were associated with spatial coordinates as well as features that were independent of spatial coordinates. We generated an input data matrix **X** containing *p* = 20 features and *n* = 500 samples, alongside *q* = 2-dimensional spatial coordinates in Cartesian space corresponding to a rough doughnut shape (Figure 2a, left column, Methods). We split features so that some features were 1) uncorrelated with spatial coordinates, 2) some features were correlated with the radial position of the sample, and 3) some features were correlated with the angular position of the samples, all with varying degrees of total noise. We then performed other common approaches for supervised or unsupervised dimensionality reduction approaches, including GraphPCA (Yang et al., 2024), PLS, and PCA. We found that in this simulation, the corresponding scores generated by the first and second wSIR’s directions were highly correlated with the angle and radius, respectively, where in contrast, GraphPCA and PLS were only able to partially recover the spatial pattern. As expected, PCA did not capture the spatial pattern and rather captured the non-spatial variables that were simulated to have the highest amount of variation (Figure 2a). Since the spatial patterns correspond to angle and radius, we conclude that wSIR is capable of prioritising the underlying linear signals in the gene expression data, which encode the non-linear relationship with the spatial positions of each cell.

**Figure 2:**
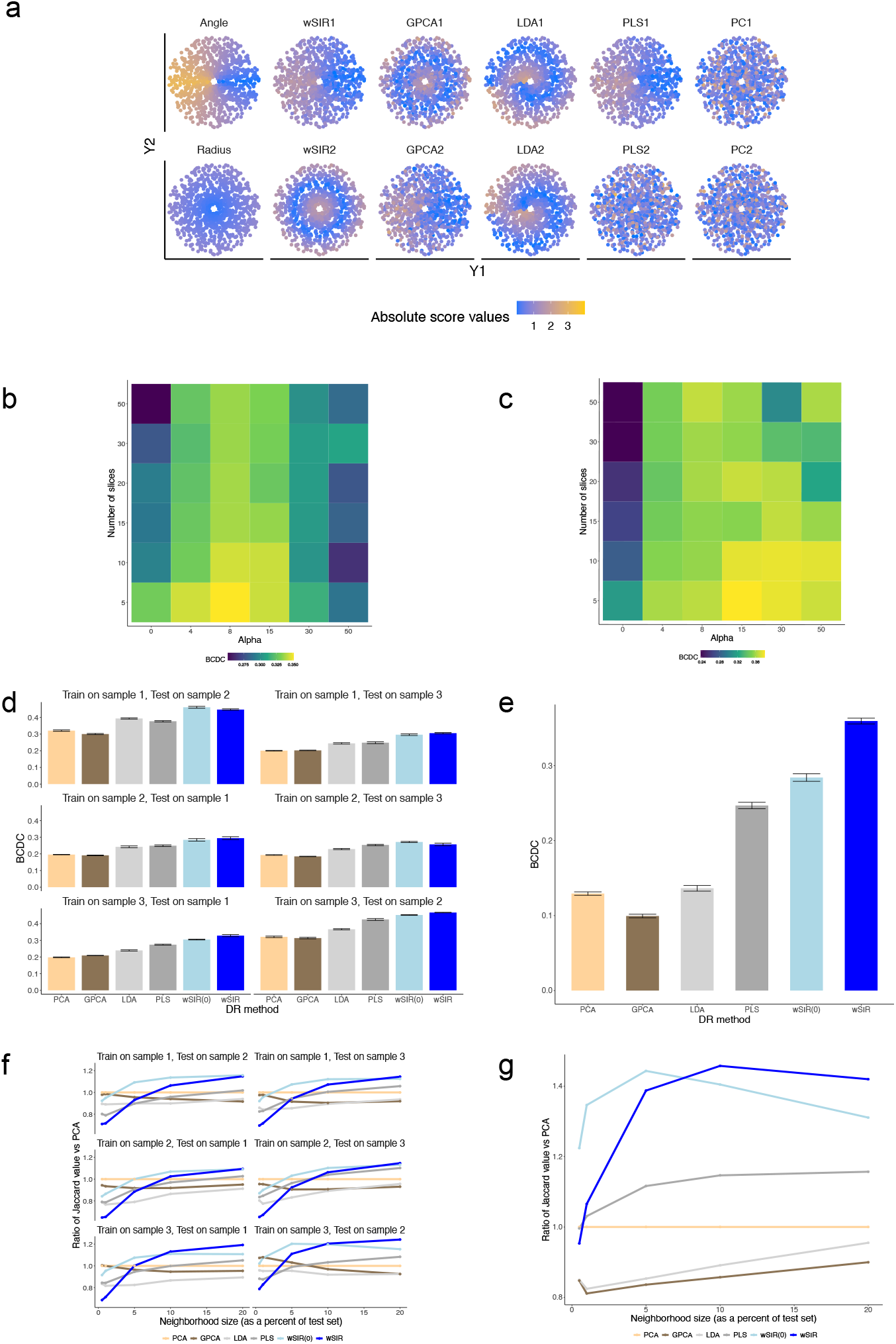
Comparison to other methods. **a**, Result of illustrative simulation where points are placed according to their simulated spatial position, and points are coloured according to the true simulated value (first column), subsequent columns are arranged according to: wSIR, GraphPCA, linear discriminant analysis (LDA), partial least squares (PLS), and PCA. The first row indicates the first direction or component selected for each method, and the second row indicates the second direction or component. **b**, Heatmap displaying the bias corrected distance correlation (BCDC) for the mouse embryo data with different choices of number of slices and slice-slice weighting alpha. **c**, similar to panel b but for the Xenium breast cancer dataset. **d**, Barplots displaying the BCDC of different dimension reduction (DR) methods for a leave-one-dataset-out evaluation of the three samples of the mouse embryo dataset. wSIR(0) indicates *α* = 0, equivalent to SIR. Error bars indicate bootstrap selection of the training set. **e**, Similar to panel d for Xenium breast cancer dataset. **f**, Line plots indicating the ability to preserve local neighbourhoods for each method, relative to PCA, using the same evaluation strategy as panel d. **g**, Similar to panel f for Xenium breast cancer dataset.

### 2.3 wSIR is robust to parameter choice

Next, we sought to examine the impact of the two main parameters in the wSIR algorithm on the performance of the corresponding low-dimensional embeddings. The main tuning parameters of interest are 1) the number of tiles to partition the spatial coordinates, and 2) Alpha (*α*), representing the scaling factor applied to the slice-slice correlation matrix (Methods). We note that for certain selections of these tuning parameters, the wSIR approach becomes equivalent to SIR (Alpha chosen as 0) and PCA (i.e. when the number of slices approaches the total number of observations in the dataset). For this exploration, we used two case study datasets from single-cell resolved transcriptomic technologies in mouse development and human breast cancer stemming from seqFISH and 10x Genomics’ Xenium platform respectively (Lohoff et al., 2021; Janesick et al., 2023) (Methods). For each combination of these tuning parameters, the quality of the low-dimensional embedding was measured by the bias-corrected distance correlation (BCDC, Methods) between it and the true spatial coordinates. The resulting heatmaps for each dataset (Figure 2b-c) show that, overall, wSIR obtains a higher BCDC than SIR, and that a high value of BCDC is maintained for parameter choices deviating from the “optimal” choices. This indicates that the performance of wSIR is relatively robust to the choice of parameters, and that a selection of a sub-optimal combination of parameters is likely to still yield useful results.

### 2.4 wSIR embeddings better preserve local and global cell-cell distances

Following our exploration of tuning parameter selection for wSIR, we examined the relative performance of wSIR compared to other dimensionality-reduction approaches used in single-cell and spatial transcriptomics analysis. Using the two case-study datasets (mouse development and human breast cancer), we performed a cross-validation study to compare the BCDC for PCA, GraphPCA, LDA, PLS, SIR (i.e. wSIR with Alpha = 0), and wSIR (Methods). In particular, we aimed to map *non-spatial* gene expression data to a spatially-informed low-dimensional embedding, so we only used spatial coordinates of the query cells to assess accuracy after projecting the cells. Given this aim, we compared wSIR with the methods that have abilities to project new gene expression data to such a low-dimensional embedding. As such, other spatially-aware dimensionality reduction approaches, such as SpatialPCA (Shang and Zhou, 2022), are not included in this comparison.

In the case of the mouse development, we performed cross-validation by selecting two of the three biological samples as reference and query, and repeating for each of the six combinations. For the human breast cancer data, we repeatedly randomly selected subsets of the cells to treat as reference and took the remainder of cells as query. Following this, we found that wSIR consistently resulted in higher BCDC values, indicating its ability to preserve spatially relevant patterns from gene expression data (Figure 2d,e). When we examined an additional metric of neighbourhood preservation (Methods), by taking the Jaccard index of local neighbourhoods of cells according to physical space and the low-dimensional embeddings, we found that, overall, either wSIR or SIR were best able to faithfully recapitulate the local neighbourhoods of cells, especially as the sizes of neighbourhoods increased as a percentage of the test sets (Figure 2f,g). Taken together, this cross-validation approach shows that wSIR has a superior performance in generating low-dimensional embeddings that faithfully recapture spatial proximities and patterns relative to other existing methods.

### 2.5 Uncovering relevant spatial biology in embryonic development

To further investigate the ability of wSIR in uncovering relevant biological patterns, we performed an exploration of single-cell spatial transcriptomics data of early mouse development (Lohoff et al., 2021). This dataset contains 351 genes profiled across three biological replicates of sagittal sections of mouse embryos at the stage of early organogenesis, and was annotated with 23 distinct cell types across the entire body plan of the developing embryo. We applied wSIR to this dataset alongside PCA as a comparison (Methods), to generate low-dimensional embeddings for the spatially-resolved cells. We noticed that all cell types were distinguished with wSIR in a similar manner to PCA (Figure 3a,b), but that there were additional subdivisions of cell types such as ‘Forebrain/Midbrain/Hindbrain’ and ‘Gut tube’ compared to PCA, illustrating that wSIR is able to subdivide heterogeneous cell types.

**Figure 3:**
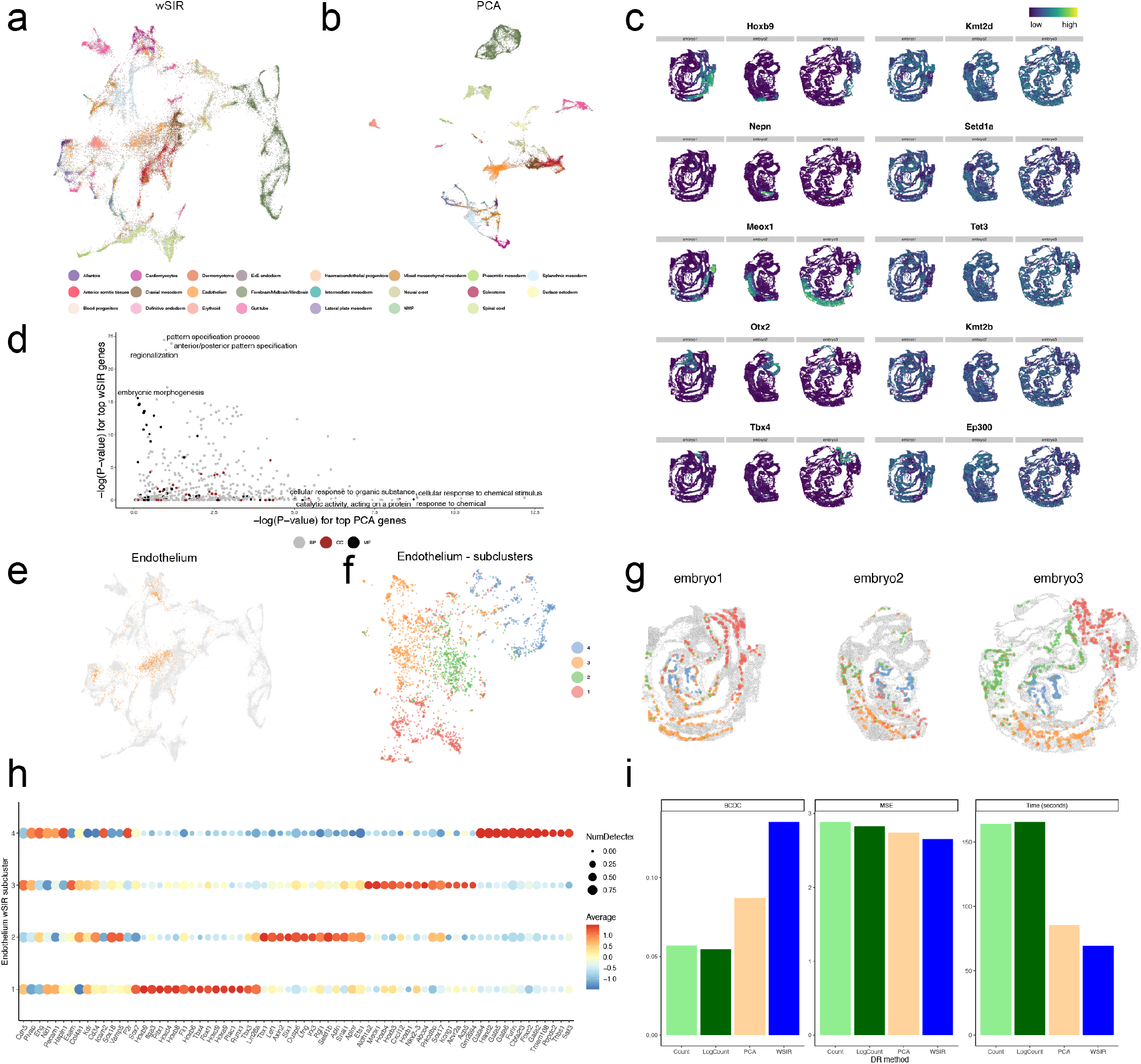
Exploration of mouse development dataset and spatial mapping benchmark with Tangram. **a**, UMAP of seqFISH mouse development cells using wSIR, coloured by original cell type annotation. **b**, Same as panel a but using PCA as underlying dimension reduction approach. **c**, Spatial expression plots of top 5 genes according to wSIR (left column) and PCA (right column). **d**, Scatterplot displaying GO terms according to − log(p − value) for enrichment of top-ranked genes in PCA (x-axis) and in wSIR (y-axis), where the most significant terms are labelled. **e**, UMAP as in panel a highlighting cells belonging to Endothelium. **f**, UMAP of Endothelium cells using wSIR, coloured by clustering of these cells. **g**, Spatial plots of Endothelium cells coloured by clusters. **h**, Expression dotplot of Endothelium clusters, the first 15 genes correspond to markers of Endothelium, and other genes correspond to specific markers for the clusters. The size of each dot represents the proportion of detected expression, and the colour indicates the average expression level relative to each group. **i**, Tangram benchmarking with the mouse development dataset. Bars correspond to each dimension reduction (DR) method (x-axis), and y-axes correspond to bias-corrected distance correlation (BCDC, left panel), mean-squared error (MSE, middle panel), and running time for Tangram in seconds (right panel).

When examining the most relevant genes according to wSIR and PCA (Methods), we identified different sets of highly ranked genes, notably, we observed genes such as Hoxb9, Nepn and Meox1 as highly ranked for wSIR. According to visual inspection, these genes are extremely localised to anatomical sites such as the tail, trunk and developing spine, respectively (Figure 3c). For the highly ranked genes in PCA, we did not observe such a marked pattern of localised expression. To examine this pattern further, we extracted the top 50 ranked genes according to wSIR and PCA, and performed functional enrichment analysis using GO terms. We found that the degree of significance for these terms were uncorrelated, and we noted relevant significant terms for wSIR including ‘pattern specification process’, ‘regionalization’, and ‘embryonic morphogenesis’, whereas the terms most significant for the PCA-ranked genes were related to epigenetic regulation and methylation enzymes and not cell type specific (Figure 3d). This indicates that wSIR prioritises gene expression variability relevant to the formation of the body plan, most clearly belonging to the formation of the head-to-tail axis.

Following the wSIR analysis of all the cells belonging to the mouse development dataset, we focused on the cells belonging to the ‘Endothelium’ cell type as we noticed these cells were especially diverse according to the wSIR low dimensional embedding (Figure 3e), with some cells appearing closer to ‘Cardiomyocytes’ and ‘Allantois’ in the UMAP representation. We subset the Endothelium cells and performed wSIR followed by graph-based clustering (Figure 3f), resulting in four identified clusters. Visualising the original locations of these cells indicates that the clusters correspond to distinct anatomical regions, such as the posterior region which includes the allantois (cluster 1) and the endocardium (cluster 4) (Figure 3g). Further examination of the marker genes associated with these clusters highlights the common expression of endothelial genes such as Cdh5, Plvap and Eng, alongside unique expression of the clusters. The posterior endothelial cluster expressed high levels of Plac1 and Tbx4, a T-box transcription factor, which plays an important role in vasculogensis within the developing allantois (Naiche and Papaioannou, 2003; Naiche et al., 2011). We also observed expression of Gata4, Gata5, and Gata6 in cluster 4 indicating the endocardium (Tremblay et al., 2018; Rivera-Feliciano et al., 2006) (Figure 3h). Taken together, our exploration highlights the utility of wSIR in providing access to spatial information during dimensionality reduction. This enables researchers to: (1) identify gene sets with important spatial roles, such as those involved in axial patterning during mouse development, and (2) uncover spatially localised cellular subtypes, such as vascular endothelial subsets arising in distinct anatomical regions. These findings underscore the potential of wSIR as a powerful tool for integrating spatial context into high-dimensional analyses, offering new insights into the spatial organisation of cellular and molecular processes.

### 2.6 Spatial prediction approaches are improved by using wSIR

Since we have shown wSIR’s ability to generate low-dimensional embeddings that are useful in uncovering spatial biology, we wondered if wSIR, as a linear dimensionality reduction approach, would be useful as an input into more complex modelling approaches. To examine this, we turned to a popular method, named Tangram (Biancalani et al., 2021), that uses deep learning to map single-cell gene expression data to spatial transcriptomics references. Tangram typically takes in a filtered gene expression matrix as input, and returns a prediction of physical locations of single cells. Using the mouse development dataset, we performed Tangram with four input types, 1) gene expression counts, 2) gene expression logcounts, 3) PCA scores, and 4) wSIR scores. To assess performance, we measured BCDC, mean-squared error, and computational runtime of the Tangram step. Overall, we found that using wSIR as the input to Tangram resulted in better performance in terms of a larger BCDC, smaller MSE, and shorter run time (Figure 3i). Overall, this investigation indicates that wSIR is a useful initial analytical step prior to applying other bespoke analyses, and is versatile in the range of downstream applications that can be employed.

### 2.7 Exploration of breast cancer spatial transcriptomics at spot- and single-cell level

Following our application of wSIR to single-cell spatial transcriptomics data, we wondered if wSIR would allow us to similarly interrogate spot-level spatial transcriptomics data to derive biologically relevant insights. The human breast cancer dataset stems from an experimental study where slices from the same tissue block were subjected also to spot-level spatial transcriptomics via the 10x Visium platform (Janesick et al., 2023). Using this spot-level data, we first subset genes to those also captured in the single-cell resolved Xenium data. We performed both wSIR and PCA, and found that we were able to recapitulate the diversity of tissue structure (Figure 4a,b), in particular identifying regions belonging to the tumour invasive region (cluster 6) and DCIS (cluster 2) (Figure 4c-e). Using the gene importance metric for wSIR and PCA (Methods), we found there was a low association between genes identified as strongly relevant by wSIR and PCA, such as MMP12 and OPRPN ranked most relevant by wSIR and PTGDS and SFRP4 ranked most relevant by PCA (Figure 4f). We noted that some of the top-ranked wSIR genes were not necessarily the most prevalently expressed (Figure 4g), for example, MMP12, a matrix metallopeptidase gene, appears to be expressed in distinct small regions of the Visium-resolved tissue, while PTGDS and SFRP4 are more prevalently expressed across the tissue. This suggests that wSIR may be able to prioritise spatial gene expression that is highly localised and potentially biologically relevant.

**Figure 4:**
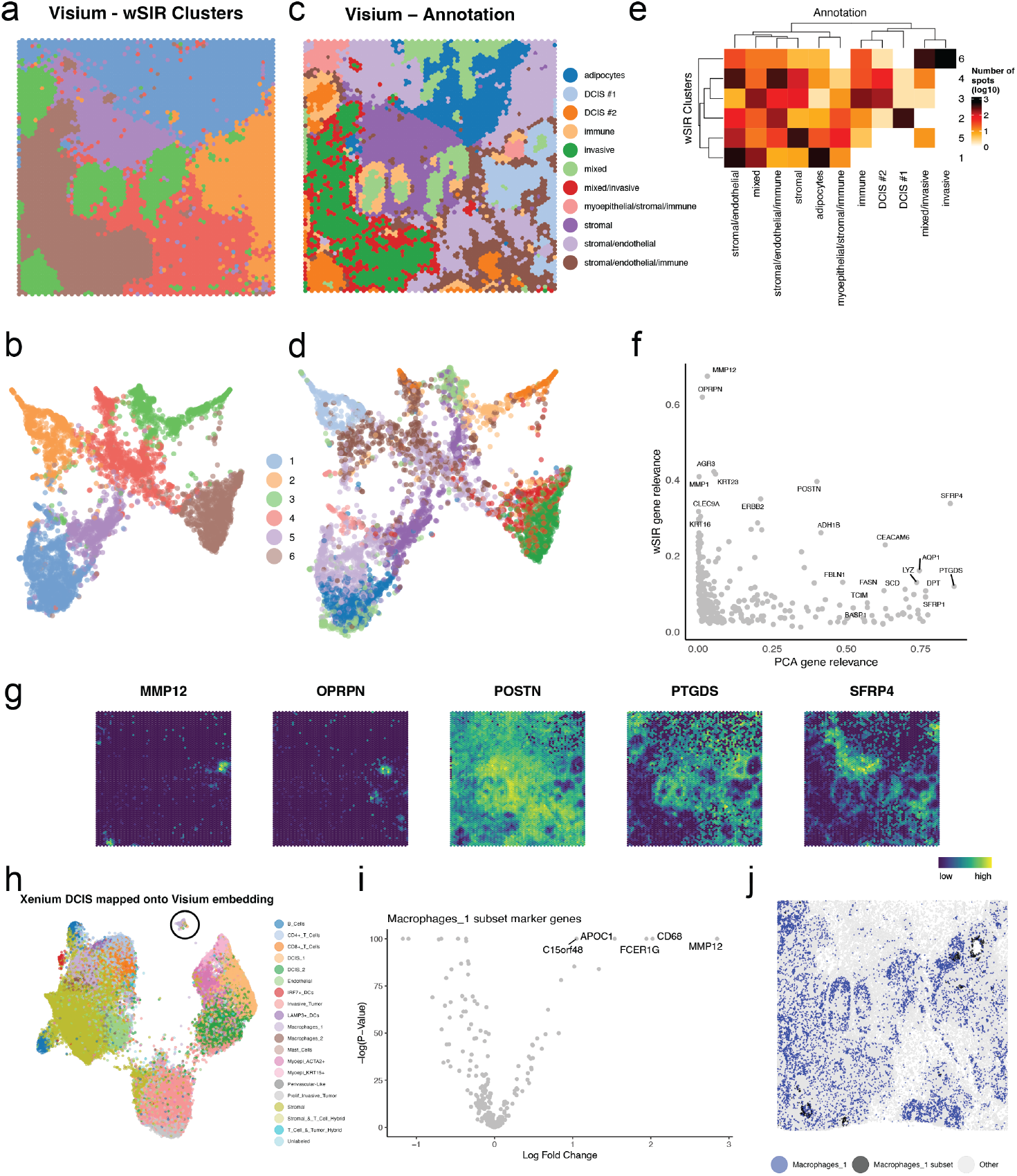
Exploration of ductal breast cancer dataset. **a**, Spatial plot of breast cancer Visium data where spots are coloured according to clustering of the wSIR dimension reduction. **b**, Corresponding UMAP plot of Visium spots with spots coloured as in panel a. **c**, Spatial plot coloured by annotation from original publication. **d**, Corresponding UMAP plot with spots coloured as in panel c. **e**, Heatmap of tabulation of number of spots belonging to wSIR clusters and original annotation. **f**, Scatterplot of gene relevance of PCA (x-axis) and wSIR (y-axis), where higher values mean more relevant genes. **h**, Spatial expression plots of selected top-ranked genes according to wSIR (MMP12, OPRPN), both wSIR and PCA (POSTN), and only PCA (PTGDS, SFRP4). **h**, UMAP of breast cancer Xenium cells according to projection to the Visium-informed wSIR low dimensional embedding. Cells belonging to a distinct group made of majority Macrophages 1 are circled in black. **i**, Volcano plot displaying differential expression testing results of the set of cells selected in panel e. **j**, Spatial plot of Xenium cells that belong to the selected group, coloured in black, corresponding to a macrophage subcluster that localise around ducts and express MMP12.

Our next question was whether we could successfully map single-cell resolved data to a low-dimensional embedding that is informed with spot-level spatial transcriptomics data. To do so, we took the Xenium-resolved human breast cancer cells, and projected the gene expression to the low dimensional space as estimated from the Visium data. We observed that the cell types as originally annotated (Janesick et al., 2023) were preserved among this projection (Figure 4h). Interestingly, we noticed that a set of cells primarily annotated as ‘Macrophages 1’ appeared as a distinct group, and this was not the case for PCA. When we selected these cells and performed differential expression analysis, we observed that the cells were marked by genes including MMP12, CD68, APOC1 and FCER1G, indicating that these cells indeed correspond to a type of macrophage that may be involved in remodelling of the extracellular matrix (Figure 4i). Since this gene expression data came from the Xenium platform, we were able to visualise their original locations (Figure 4g). In doing so, we noted distinct positioning of these cells around some of the DCIS (as annotated in the original publication), suggesting a possible role of these cells in the progression into invasive tumours. Taken together, our in-depth case study of single-cell-resolved and spot-resolved human breast cancer spatial transcriptomics data encapsulates the ability of wSIR to uncover relevant signals for spatial cancer biology.

## 3 Discussion

In this paper, we have introduced wSIR, an approach to perform spatially-aware dimensionality reduction of single-cell and spot-level spatial transcriptomics data. wSIR can handle multiple distinct biological samples, and enables mapping of non-spatial gene expression data to a low-dimensional embedding that is informed by spatial coordinates. We have implemented wSIR as an R package, with scalability due to efficient Rcpp implementation. We have shown relatively superior performance of wSIR compared to other dimensionality reduction approaches, and demonstrated wSIR’s applicability in two real-world biological contexts.

A current limitation of wSIR is that an initial step of feature selection is needed for very large datasets, e.g. whole transcriptome data. While there are works in statistics research on implementing sparsity to SIR via the Lasso (Lin et al., 2019), future work would focus on incorporating such regularisation approaches to the wSIR context in terms of both the slicing mechanisms for spatial transcriptomics data, and taking the slice-slice correlations into account. Adding further complexity to the modelling step could be met with a decrease in computational scalability, so future work would focus on ensuring that any algorithmic advance is not met with a large drop in computational efficiency.

A key advantage to wSIR is its ability to take into account multiple samples to generate a single low-dimensional loading and embedding. We currently do so by assuming that the slice-slice correlation of slices belonging to separate samples is zero. With the increase in throughput of spatial transcriptomics studies and inevitable rise in variety and complexity of the studies (Velten and Stegle, 2023), it is important to consider different levels of replication. For example, datasets can contain distinct spatial samples that are 2D slices from the same 3D tissue block across multiple individuals, and care will need to be taken to build an appropriate slice-slice correlation matrix in these situations. There are algorithms that computationally ‘register’ 2D samples into a 3D structure, but there is a potential circularity issue in using gene expression data to register spatial coordinates, then using spatial coordinates to reduce dimension of gene expression data. Future work could consider tackling these tasks jointly, potentially leading to more accurate results.

We envisage wSIR being used in a variety of contexts, especially as spatial transcriptomics data becomes more widespread. Effectively using spatial information is a challenge in contemporary spatial transcriptomics analyses, and simply using tools designed for scRNA-seq can hinder the ability to draw insight in terms of spatial biology. Therefore, we see the potential of wSIR to ensure that spatially relevant variation is not lost, and to enable more comprehensive understanding of spatial biology in several contexts.

## 4 Materials and Methods

Dimension reduction (DR) refers to statistical methods aiming to reduce the dimension, typically the number of columns, of a dataset to address the curse of dimensionality and remove noise from the data. Let **X** be an *n × p* matrix of gene expression features, where *n* is the number of observations (cells) and *p* is the number of genes, where *p* is typically large. Dimension reduction methods aim to find a low– dimensional score *n × d* matrix **V**, with *d*≪ *p*, where different methods differ in what information of **X** (e.g total variances, graph structures) need to be retained as much as possible in **V**.

In our spatial transcriptomic context, in addition to **X**, we have an *n× q* matrix of response **Y** containing spatial coordinates of cells with *q* = 2 or *q* = 3. Therefore, we aim to find such a matrix **V** so that **V** preserves the ability of **X** to predict **Y** (Li, 2018), i.e **Y** ⊥ **X** | **V** with ⊥denoting statistical independence and | denoting conditioning. This is precisely the aim of “sufficient dimension reduction”, which is an active research area that gains increasing popularity among practitioners. Among sufficient dimension reduction methods, SIR is the most widely used due to its generality, simplicity, and computational efficiency. Since its first introduction in Li (1991), SIR has been adapted into many complex settings, including missing data (Li and Lu, 2008; Xiao and Zhang, 2022) and high-dimensional features with *n* ≪ *p* (Lin et al., 2019; Li et al., 2023). Nevertheless, most recent works on SIR assume a univariate response, i.e. *q* = 1. While a few works have extended SIR to the multivariate response settings (Li et al., 2003; Coudret et al., 2014), our paper is the first to propose SIR for responses with spatial coordinates. Without the loss of generality, we assume each column of the feature matrix **X** is centered.

### 4.1 A brief review of sliced inverse regression

As a linear sufficient dimension reduction method, SIR assumes **V** = **XB** where **B** is a *p× d* loading matrix **B** such that **Y**⊥ **X** |**V**. This conditional independence relationship is invariant for any rotation of **V** by an orthonormal *d × d* matrix **C**, i.e **Y** ⊥ **X**| **V** is equivalent to **Y** ⊥ **X** | (**VC**), so the loading matrix **B** is only identifiable up to rotation. Compared to non-linear sufficient dimension reduction methods, linear dimension reduction methods are more interpretable and scalable, and they enable a straightforward projection of a new dataset into the lower-dimensional space.

The basic algorithm for SIR is as follows. Let 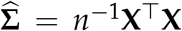 be the sample covariance matrix of the gene expression data and 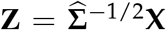 be the standardised covariates matrix. In the first step, all of the observations are allocated into one of the *H* slices, where each slice has observations with similar value for the response variables *Y*, where *H* is a pre-specified user input. Let **z**_*h*_ be a *p ×* 1 column vector consisting of the average of all the standardised covariates within the *h*th slice for *h* = 1, …, *H*. In the second step, let 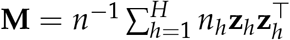 be a sample covariance matrix of these sliced means, where *n*_*h*_ is the number of observations within the *h*th slice. Then, an eigendecomposition is performed on **M** to obtain its eigenvectors **u**_1_, …, **u**_*d*_, and each column of the final SIR estimates of the loading matrix 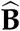 is obtained as 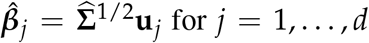 for *j* = 1, …, *d*. We note that while any two eigenvectors **u**_*j*_ and **u**_*k*_ are orthogonal to each other for *j*≠ *k*, the columns 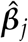 and 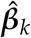 are not. The column space of **B** is finally estimated by the column space of 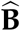.

There are two main challenges of adapting the above basic SIR algorithm to our settings with spatial coordinates responses. First, it is not clear how to form slices, second, each slice in our setting corresponds to one spatial location, so these slices are spatially correlated with each other. Nevertheless, in the basic SIR algorithm, this spatial information is ignored after the slices are formed. To overcome these challenges, we introduce a general weight matrix into the computation of the sample covariance matrix of the sliced mean, where the off-diagonal elements are explicitly used to account for the correlation among the slices.

#### Slicing methods for spatial coordinates responses

To form slices of the data with a multivariate response, for each sample, we assign cells to *H* = *k*^2^ tiles, where *k* is the number of slices representing the number of non-overlapping bins of equal length along each axis direction in the 2D coordinate. In general, the number of cells/spots within each slice should be large enough to benefit from the SIR mechanism, noting that in the extreme case, if each cell/spot is within its own slice then the approach is akin to PCA. While there are practical discussions around slicing in the SIR literature (Girard et al., 2022) and general heuristics can be followed (e.g. at least 25 cells), the number of slices is treated as a tuning parameter and a selection is made according to the largest cross-validated BCDC.

### 4.2 wSIR: Spatial weighting matrix to handle correlations among the slices

In the second step of the basic SIR algorithm, we can write the matrix **M** in the form

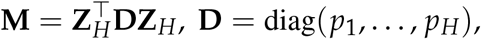

where **Z**_*H*_ is the *H× p* matrix whose *h*th row is the vector of average of the standardised covariates and *p*_*h*_ = *n*_*h*_/*n* for *h* = 1, …, *H*, and *H* is the number of tiles. Since the matrix D is diagonal, no correlation among the slices are accounted for in this computation of **M**. To allow for the correlation among the slices and preserve the proportion of observations within these slices, we propose to compute **M** to be

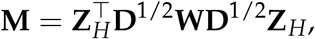

where **W** is a *H × H* correlation matrix, whose diagonal elements are one and off-diagonal elements are between 0 (inclusive) and 1. Since these tiles correspond to spatial locations, the higher proximity between two tiles, the more correlated they are. Hence, we construct the off-diagonal elements *w*_*ij*_ of **W** from the distance between the centroid of the *i*th and *j*th tile as follows.

Let *S*_*h*_ be the index set for the *h*th tile, i.e *S*_*h*_ = {*i* : *ith* observation belong to the *h*th tile} for *h* = 1, …, *H*. Then the spatial coordinates for the centroid of the *h*th tile is

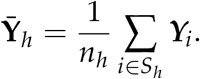

Therefore, the Euclidean distance between the centroid of the two tiles (*i, j*) is 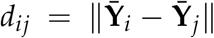 for *i, j* = 1, …, *H*. We then compute

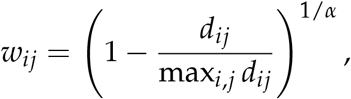

where *α >* 0 is another tuning parameter. This choice of *w*_*ij*_ in the equation above ensures that all the resulting *w*_*ij*_ are between 0 and 1, and a higher distance *d*_*ij*_ leads to a lower value for *w*_*ij*_ as desired. Furthermore, we include the *α* to ensure our resulting method can include the basic SIR as a special case; indeed the basic SIR that ignores all the correlations among the slices correspond to *α* = 0.

#### Selecting the number of wSIR directions

To select the number of directions *d* for the low-dimensional score, we employ a cumulative variance threshold when performing the eigendecomposition of the matrix *M*. Particularly, let 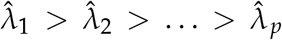 be the eigenvalues of **M**, so each 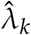 is the variance of the score **Z**_*H*_**u**_*k*_, for *k* = 1, …, *p*. We then choose an estimate 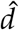 to be the minimum number of directions that explains a certain percentage of total variance across all the scores, i.e

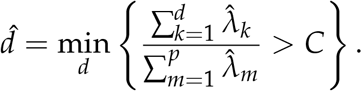

As a default, we look for the minimum number of directions that captures at least 99% of the variance, so *C* = 0.99. Otherwise, as often done in PCA, we can keep the number of wSIR directions to be 50, particularly when we use wSIR embeddings for other downstream analyses (such as clustering or prediction).

#### Selecting tuning parameters for Wsir

For these other parameters in the wSIR algorithm, including the number of tiles *H* and the coefficient *α* in the computation of the weight matrix, we use a 10-fold cross-validation. Specifically, we split our data into a training set (**X**_train_, **Y**_train_) and a validation set (**X**_val_, **Y**_val_). For each potential candidate of *H* and *α*, we computed the corresponding wSIR loading 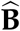 in the training data with the number of components being selected from the above cumulative variance threshold. Then, we computed the score on the test set 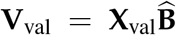, and we choose the combination of (*H, α*) that maximises the distance correlation between the two matrices **Y**_val_ and **V**_val_ (Székely et al., 2007). This distance correlation metric is widely used in the statistics literature to measure the dependence between two datasets, encompassing both linear and non-linear relationship among them (Sheng and Yin, 2016; Nghiem et al., 2022; Zeng et al., 2024). We outlined the computation of this metric below.

#### Applying wSIR to multiple samples

When we use multiple samples in the training set for a dimension reduction procedure, we must account for possible differences between both the gene expression data and the spatial coordinate systems (e.g coordinate (0,0) in one sample may not correspond to coordinate (0,0) in another sample). These concerns can both be addressed by the weight matrix. In the weight matrix, a value of 0 corresponds to no spatial correlation between the tiles. For a pair of tiles from different samples, we set the corresponding entry in the weight matrix to 0.

The weight matrix has a dimension 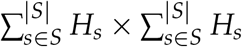, where *S* is the set of all samples, *s* is one of the samples, and *H*_*s*_ is the number of tiles in sample. For any entry *ω*_*ij*_ of the weight matrix **W**, it will take value 0 if tiles *i* and *j* come from different samples. If they are from the same sample, its value will be calculated in the usual way as if there was only one sample in the wSIR computation. As a result, we are not assuming any spatial correlation between tiles from different samples.

#### Ranking variables by importance in wSIR

Following the fitting of wSIR, we examine the estimated loadings matrix 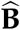 to obtain a ranking of the variables that are most relevant in the dimensionality reduction. Recall that the matrix **B** is only identifiable up to an orthogonal rotation, but the ranking of variables within each column of **B** is generally not invariant to such a rotation. Hence, we remove the effect of rotations by computing the corresponding *p × p* projection matrix 𝒫(**B**) = **B**(**B**^⊤^**B**)^−1^**B**^⊤^. Then, a measure of importance for the *j*th gene expression feature is the corresponding diagonal element of 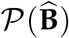, i.e 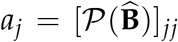 for *j* = 1, …, *p*. The values of *a*_*j*_ range from 0 to 1, with 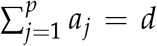 a higher value of *a*_*j*_ indicates the *j*th feature is more important. Intuitively, a high value of *a*_*j*_ indicates a relatively large loading of the *j*th gene expression feature in *at least* one of the *d* dimensions; a low value of *a*_*j*_ indicates the corresponding gene expression has a small loading in *all* dimensions. This measure of variable importance is commonly used in sufficient dimension reduction, see for example Tan et al. (2018); Nghiem and Hui (2024).

We then rank the variables in descending order according to these scores. The user may select a set number of top-ranked variables as relevant, e.g. the top 50 genes. We note that this approach to ranking variables is also applied to the loadings estimated using PCA.

#### Projecting non-spatial data using wSIR

Since a fitted wSIR model only involves the feature-level information of spatial omics data, non-spatial data can be projected into the spatially-aware low-dimensional space as estimated from spatial data. This is done by employing the same projection as outlined in the subsection 4.2 below, but for centred and scaled non-spatial gene expression data, 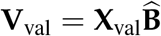.

### 4.3 Comparison with existing methods

We compare wSIR with PCA, LDA, PLS (Abdi, 2010), and GraphPCA (GPCA) (Yang et al., 2024). PCA forms the low-dimensional scores **Z** that preserve as much as total variability in **X**; as a common practice, we kept the number of PCs to be 50 (Kobak and Berens, 2019). For LDA, we create a categorical response variable with *H* categories, each corresponding to one slice. LDA finds *H* − 1 linear combinations of **X** that separate these categories. PLS finds the components that maximise the sample covariance between **X** and the spatial coordinates **Y**; we selected the number of low-dimensional components by cross-validation as implemented in the `plsr` function of the `pls` R package (Liland et al., 2024). GPCA is a regularised version of PCA; it aims to construct low-dimensional scores **Z** of **X** that minimise the reconstruction error while preserving the graph structures among the cells. GPCA requires an *n × n* cell-cell adjacency matrix, which is obtained from computing *k*-nearest neighbour graph on the spatial coordinates **Y**. As implemented in the `GraphPCA` Python package, the default choice is *k* = 5 for ST platform, and the regularisation parameter for the graph constraint is fixed at 0.5.

## 5 Datasets and case studies used in this study

### 5.1 Synthetic data

We generated a simulated dataset where we know the true low-dimensional spaces (ground-truth) and used them to demonstrate wSIR and compare it with other popular dimension reduction methods. Specifically, we generated the spatial coordinates **Y**_*i*_ = (*Y*_*i*1_, *Y*_*i*2_) from *Y*_*i*1_ = *R*_*i*_ cos(*θ*_*i*_) and *Y*_*i*2_ = *R*_*i*_ sin(*θ*_*i*_), where the radius *R*_*i*_ and *θ*_*i*_ are independently random numbers drawn from the Uniform(0.1, 1) distribution and the Uniform(− *π, π*) distribution for *i* = 1, …, *n*. The *n × p* gene expression feature matrix **X** is decomposed into three blocks **X** = [**X**_1_, **X**_2_, **X**_3_], where **X**_1_ is an *n × p*_1_ matrix whose elements are independently generated from *N*(0, 5) distribution, **X**_2_ is an *n× p*_2_ matrix whose each element in the *i*th row are drawn from *N*(*R*_*i*_, 0.5) distribution, and **X**_3_ is an *n× p*_3_ matrix whose each element in the *i*th row is independently drawn from *N*(*θ*_*i*_, 0.5) distribution for *i* = 1, …, *n*. We set *n* = 1000, *p*_1_ = 10, *p*_2_ = *p*_3_ = 5, so *p* = *p*_1_ + *p*_2_ + *p*_3_ = 20. In this synthetic data, the ground-truth two-dimensional low-dimensional scores that fully capture the spatial information are **V** = (**X**_2_, **X**_3_) = **XB**, where **B** = [**B**_1_, **B**_2_] with 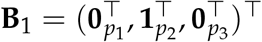 and **B**_2_ = (**0**_*p*1+*p*2_, **1**_p3_)^T^., where **0**_*k*_ and **1**_*q*_ denote a *k ×* 1 and *q×* 1 vectors of zero and one, respectively. Figure 2 (panel a, first column) plots a generated sample of spatial coordinates (*Y*_1*i*_, *Y*_2*i*_), coloured by the magnitude of *R*_*i*_ (top) and *θ*_*i*_ (bottom) for *i* = 1, …, *n*.

### 5.2 Human breast cancer dataset

We downloaded single-cell spatial transcriptomic data for human breast cancer from the 10x Genomics website (see Data Availability). We retained only spatial coordinates and gene expression belonging to replicate 1. We performed filtering of low-quality cells and filtering of lowly expressed genes, prior to performing a logcount normalisation on the retained cells. We used the cell annotation provided in the publication, resulting in a dataset with *n* = 163, 998 cells and *p* = 313 genes. To obtain PCA and wSIR results for this dataset, we performed PCA (50 PCs) using the `runPCA` function in the `scater` R package, and we performed wSIR with *H* = 30^2^ tiles and*α* = 4. When comparing the performance of the wSIR with other methods in dimensionality reduction alone, (such as the results in Figure 2), we used the proportion of explained variance threshold *C* = 0.99 to determine the number of wSIR dimensions. When wSIR low-dimensional embeddings were used for further downstream analyses, we kept the number of dimensions to be 50. To facilitate visualisation, for both PCA and wSIR, we performed UMAP using the `runUMAP` function in the `scater` R package. We identified the top 50 most relevant genes according to the procedure outlined above, and performed functional enrichment analysis using Gene Ontology (GO) using the `clusterProfiler` R package, retaining GO terms containing between 20 and 200 genes, and set all the genes appearing in the dataset as the gene universe. We performed differential expression analysis using the `findMarkers` function in the `scater` R package.

We also downloaded the corresponding spot-level 10x Genomics’ Visium dataset from the same website. We performed wSIR using the same parameters except with *H* = 20^2^ tiles and a maximum of 30 directions. We projected the single-cell Xenium data into the Visium-resolved low-dimensional embedding using the `projectWSIR` function in our `wSIR` R package. We performed differential expression analysis of the macrophage subset using the `findMarkers` function in the `scater` R package.

### 5.3 Mouse development dataset

We downloaded single-cell spatial transcriptomic data from a recent study of early organogenesis (Lohoff et al., 2021) using `MouseGastrulationData` Bioconductor package. This dataset comprises three biological replicates of sagittal sections of mouse embryos, totalling *n* = 57, 536 cells measured across *p* = 351 genes. We removed the gene *Xist* from further analysis, since the embryos are a mix of male and female, and filtered cells annotated as ‘Low quality’ as well as cells containing fewer than 10 counts and fewer than five unique genes. We performed normalisation using the `logNormCounts` function in the `scater` R package. We performed wSIR of the spatial gene expression using *H* = 30^2^ slices, *α* = 4, where the number of dimensions were selected as in Section 5.2. Further visualisation using UMAP and GO enrichment testing was the same as for the human breast cancer data. For the endothelium data, we used the same wSIR parameters, and performed clustering of the cells using the `bluster` R package with infomap graph-based clustering.

## 6 Evaluation metrics

### 6.1 Distance correlation

First, we are interested in the overall association between the spatial coordinates and the low-dimensional scores, as measured by the bias-corrected distance correlation (BCDC). Given the spatial coordinates **Y** and a low-dimensional score matrix **V**, each with *n* rows, let **Y**_*i*_ and **V**_*i*_ be the *i*th row of **Y** and **V**, respectively. Let *A*_*ij*_ and *B*_*ij*_ be the pairwise distance between the *i*th and *j*th row of **Y** and **V**, i.e *A*_*ij*_ = ∥**Y**_*i*_ − **Y**_*j*_∥ and *B*_*ij*_ = ∥**V**_*i*_ − **V**_*j*_∥ for *i, j* = 1, …, *n*. The centred distances on **Y** are given by

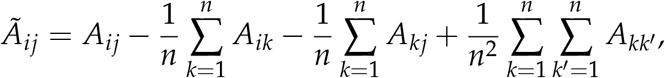

and similar definition holds for 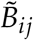. Hence the bias-corrected distance correlation (BCDC) is defined to be

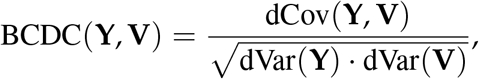

with the bias-corrected distance covariance

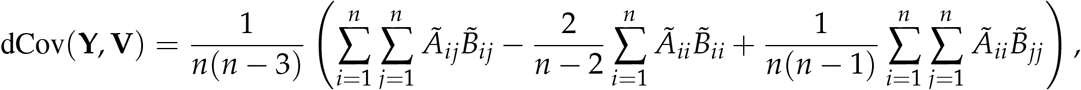

and the bias-corrected distance variances dVar(**Y**) = dCov(**Y, Y**) and dVar(**V**) = dCov(**V, V**). A higher BCDC implies a strong association between **Y** and **V**, encompassing both linear and non-linear relationship.

### 6.2 Average Jaccard coefficient

We also evaluate the performance of dimension reduction methods in how well they preserve the neighbourhood structure among the cells. Particularly, let 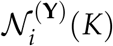 be the index set of the *K* nearest neighbours for the *i*th observation from the spatial coordinates matrix **Y**, i.e

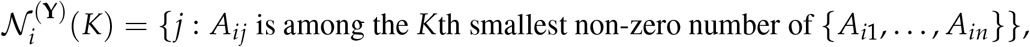

and similar definition holds for 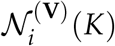 with *A*_*ij*_ replaced by *B*_*ij*_. The Jaccard index for the *i*th observations measures the similarity between the two above sets, i.e

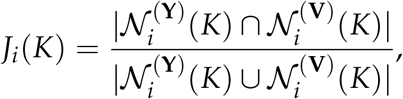

where ∩, ∪ denote the set intersection and union, respectively, and | *·* | measures the number of elements in the set. For each value of *K*, we compute the average Jaccard value 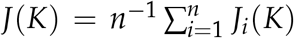; in our results, we select different values of *K* corresponding to (0.5, 1, 5, 10, 20)% the number of observations in a test set.

### 6.3 As input to Tangram spatial mapping

We used Tangram, a deep learning model for alignment, to explore the applicability of wSIR dimensions in downstream analysis. Tangram maps a query *n*_*sc*_ *× g* single cell gene expression matrix to a reference *n*_*sp*_*× g* spatial location gene expression matrix, where *n*_*sc*_, *n*_*sp*_ and *g* are the number of single cells, the number of spatial spots, and genes used for training, respectively. This mapping is characterised by a *n*_*sp*_ *× n*_*sc*_ matrix **Q** where each entry *q*_*ij*_ is the probability that the *i*th single cell being in the *j*th spatial spot, so 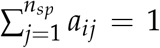 for *i* = 1, … *n*_*sc*_. The mapping is performed iteratively under a minimising loss function, according to which the cosine similarity between the gene expressions of the mapped single cells and reference spatial data cells should be as high as possible. We applied Tangram to two embryo replicates from the mouse embryo dataset, by treating replicate 2 as the query single cell dataset, i.e *n*_*sc*_ = *n*_2_ and ignoring the spatial locations, and treating replicate 1 as the reference spatial dataset, i.e. *n*_*sp*_ = *n*_1_, where *n*_1_ and *n*_2_ are the number of cells on the replicates 1 and 2, respectively. To predict the location for the query single cell data, we adopted the approach used in Li et al. (2023), which assigns the *i*th query single cell to the *j*th spatial location with the highest probability value. We applied this workflow to low-dimensional embeddings generated by wSIR obtained as outlined in Section 5.3 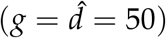, from which we predicted spatial coordinates for query single-cell samples (i.e. embryo 2), and compare the predicted coordinates with the actual spatial coordinates in that embryo. We compared the performance of this approach (wSIR + Tangram) versus (a) the same workflow with input being PCA scores of the same dimensions (PCA + Tangram), and (b) two naïve-Tangram analyses with raw (Count) and normalised (LogCount) gene expression (*g* = *p* = 351).

## Data Availability

This study used publicly available data. The mouse embryo seqFISH data was accesssed via Bioconductor (version 3.19) ExperimentHub package MouseGastrulationData (version 1.18.0). The breast cancer 10x Genomics Xenium data was accessed via the 10x Genomics website (https://www.10xgenomics.com/products/xenium-in-situ/preview-dataset-human-breast) on 2 February 2024 and processed using `MoleculeExperiment` (version 1.5.1). The matching Visium data was accessed by the same website on 1 December 2024 and processed using `DropletUtils` (version 1.24.0).

## Code Availability

All analyses were performed in R (version 4.4.2). The wSIR software is available as an R package at https://github.com/SydneyBioX/wSIR and has been submitted to the Bioconductor Project. Scripts for analysis and figure panels in this manuscript are available at https://github.com/SydneyBioX/wSIR2024.

## Acknowledgements

We thank our colleagues in the University of Sydney School of Mathematics and Statistics, Charles Perkins Centre, and Sydney Precision Data Science Centre for their support and intellectual engagement. The following sources of funding are gratefully acknowledged. S.G. was supported by an Australian Research Council DECRA Fellowship (DE220100964). P.P. was supported by the Chan Zuckerberg Initiative DAF, an advised fund of Silicon Valley Community Foundation (2022-249319 to S.G.). L.T.G.H. was supported by a Wellcome Early-Career Award (226309/Z/22/Z). B.G. was supported by Wellcome (206328/Z/17/Z and 203151/Z/16/Z) and UKRI Medical Research Council (MC PC 17230). A.S. is supported by the generosity of Deborah and John McMurtrie and the Petre Foundation and grants from the Breast Cancer Research Foundation (BCRF-23-209), NHMRC (APP2018440) & NBCF (IIRS-23-074). The funding sources mentioned above had no role in the study design; in the collection, analysis and interpretation of data, in the writing of the manuscript and in the decision to submit the manuscript for publication.

The authors declare no competing interests.

## Author Contributions

L.H.N. and S.G. conceived the study. M.W. developed the method and software and performed data analysis with supervision from L.H.N. and S.G.. P.P. performed the spatial mapping benchmarking. S.G. and L.T.G.H. interpreted the development results with input from B.G. J.M., H.Z. interpreted the breast cancer results with input from A.S. and S.G.. M.W., L.H.N. and S.G. wrote the manuscript. All authors read and approved the final version of the manuscript.

